# RECUR: Identifying recurrent amino acid substitutions from multiple sequence alignments

**DOI:** 10.1101/2025.04.29.651261

**Authors:** Elizabeth HJ Robbins, Yi Liu, Steven Kelly

## Abstract

Identifying recurrent changes in biological sequences is important to multiple aspects of biological research -from understanding the molecular basis of convergent phenotypes, to pinpointing the causative sequence changes that give rise to antibiotic resistance and disease. Here, we present RECUR, a method for identifying recurrent amino acid substitutions from multiple sequence alignments that is fast, easy to use, and scalable to thousands of sequences. We demonstrate the utility and performance characteristics of RECUR on a data set of surface glycoprotein (S) protein sequences from SARS-CoV-2 – identifying widespread recurrent evolution throughout the protein. Structural analysis of the recurrently evolving sites revealed significant enrichment in the exposed receptor-binding S1 subunit and at the interface with the human angiotensin-converting enzyme 2 (hACE2), whereas recurrent substitutions were depleted at the trimeric interface of the S protein. Finally, *in silico* modelling showed that recurrent substitutions have primarily acted to stabilise the trimeric interface, but had no consistent effect at the hACE2 interface, suggesting that evolution at these sites has been shaped by opposing selection pressures – balancing the need to maintain or enhance hACE2 binding with pressures to diversify and evade host immune responses. A standalone implementation of the algorithm is available under the GPLv3 licence at https://github.com/OrthoFinder/RECUR.

## Introduction

Recurrent evolution arises when the same biological innovation evolves independently on multiple occasions. It is found across the tree of life and can be observed at the phenotypic and genotypic level (Stern 2013). At the phenotypic level, recurrent evolution can provide insight into the role that natural selection plays in overcoming environmental constraints. Paradigm examples of recurrent phenotypic evolution include the independent evolution of wings in pterosaurs, insects, bats, and birds (Hunter 2007); the independent evolution of high-resolution camera-like eyes in vertebrates, cephalopods, and arthropods (Nilsson 2013); and the independent evolution of the C_4_ carbon concentrating mechanism in plants (Sage 2004). At the genotypic level, recurrent evolution can help uncover the mechanistic basis of disease (Ingram 1957; Cutting, et al. 1990; Davies, et al. 2002; Olivier, et al. 2010; Jänne, et al. 2022), how proteins change to better suit environmental conditions (Bull, et al. 1997; Christin, et al. 2007; Christin, et al. 2008; Yokoyama, et al. 2008; Christin, et al. 2009; Liu, et al. 2010; Projecto-Garcia, et al. 2013; van Ditmarsch, et al. 2013; He, et al. 2021; Robbins and Kelly 2024), and how organisms defend against toxic molecules (including antimicrobials, antivirals, herbicides and pesticides) (Dobler, et al. 2012; Feldman, et al. 2012; Toprak, et al. 2012; Zhen, et al. 2012; Brodie III and Brodie Jr 2015; Ujvari, et al. 2015; Iketani, et al. 2023). Collectively, substantial insights into multiple biological phenomena at disparate scales can been gained by studying recurrent evolution.

In order to identify recurrent molecular evolution, biological sequences need to be analysed in the context of evolutionary history (Figure 1). Phylogenetic analysis of sequence change allows the reconstruction of ancestral sequences (the internal nodes in the phylogenetic tree), enabling the direction and frequency of sequence change to be determined. Convergent and parallel evolution are both forms of recurrent evolution that can be inferred through such phylogenetic analyses. Convergent evolution refers to the independent acquisition of a trait from dissimilar ancestral traits, e.g., X → Y and Z → Y. Meanwhile, parallel evolution requires both the ancestral and descendent traits to be identical, e.g., multiple independent instances of X → Y. Here, the identity of the sequence change – whether arising from identical or non-identical ancestral states – is important and is independent of the rate at which the change occurs. In contrast, other measures of molecular adaptation, such as tests for positive selection quantify the rate of sequence change, specifically the ratio of the rate of non-synonymous to synonymous substitution (*d*_N_/*d*_S_), without considering the specific types of substitutions occurring. Thus, recurrent evolution provides a complementary perspective to rate-based approaches for studying molecular adaptation, focusing not just on how rapidly sequence change occurs, but on whether similar evolutionary outcomes are repeatedly achieved across the phylogeny.

**Figure 1.**
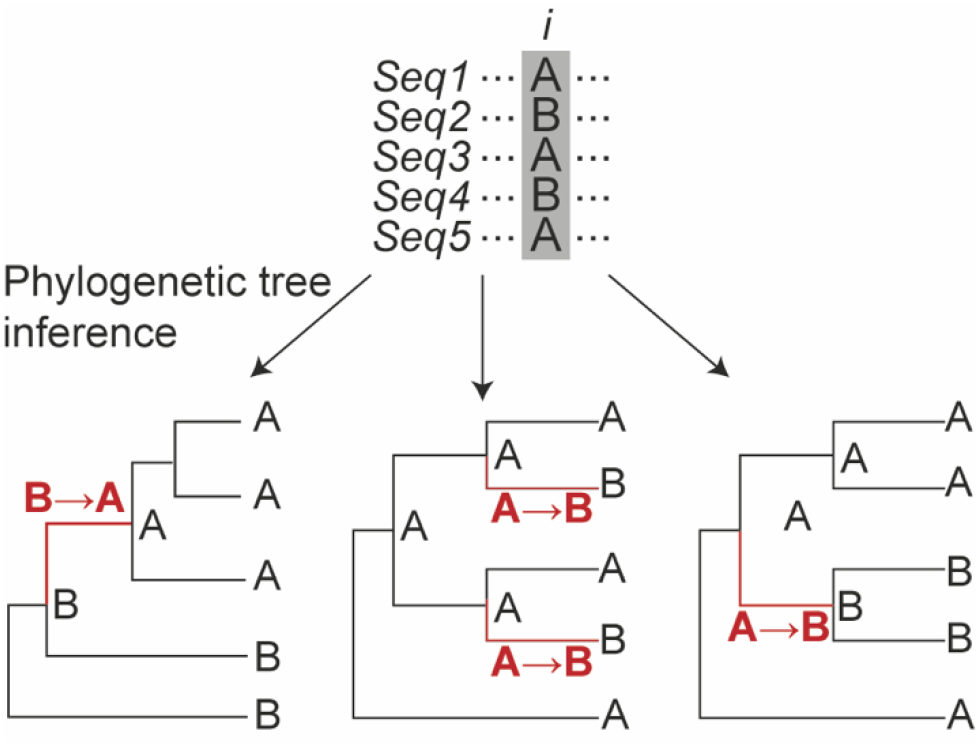
The evolutionary context of sequences in a multiple sequence alignment (MSA) dictates the substitutions inferred. A single site *i* from a MSA is shown on top with three alternative phylogenetic tree topologies below. Depending on the topology, different ancestral states are reconstructed at the internal nodes, leading to variation in the direction and frequency of substitutions.

Despite the utility of identifying recurrent evolution to the study of molecular adaptation there are currently no tools or methods that can easily identify recurrent molecular evolution directly from a multiple sequence alignment. Here, we present RECUR, a phylogenetic tool designed to address this gap by identifying recurrent substitutions, specifically parallel substitutions, that have occurred in a protein or codon multiple sequence alignment. RECUR takes a multiple sequence alignment as input and identifies all recurrent sequence substitutions present within the evolutionary history of that alignment and their associated statistics. RECUR is fast and scalable to thousands of sequences, and we exemplify its utility on an alignment of 123,126 SARS-CoV-2 surface glycoprotein sequences.

## Results

### Workflow and overview of RECUR

RECUR was designed to identify recurrent amino acid substitutions that have arisen during the evolution of a set of protein sequences. RECUR requires as input a multiple sequence alignment of protein sequences or an equivalent codon alignment. The method returns 1) the complete set of substitutions observed in the evolutionary history of that alignment. 2) The complete set of substitutions that are recurrent including whether any reversions (from the derived state back to the ancestral state) have also taken place. 3) A statistical analysis of the recurrent substitutions to identify those that have occurred more frequently than expected given the number of sequences, their phylogenetic relationship, and the underlying model of sequence evolution.

An overview of the RECUR workflow is provided in Figure 2. In brief, a phylogenetic tree is constructed from the input multiple sequence alignment and ancestral sequences are inferred for all internal nodes of the phylogenetic tree. All amino acid substitutions are then identified by assessing changes in the protein sequence along every branch in the phylogeny. Simulated multiple sequence alignments are then generated using the inferred ancestral sequence, the inferred tree, and the best-fitting model of sequence evolution and evaluated to identify those recurrent substitutions that have occurred more frequently than expected by chance (see Methods).

**Figure 2.**
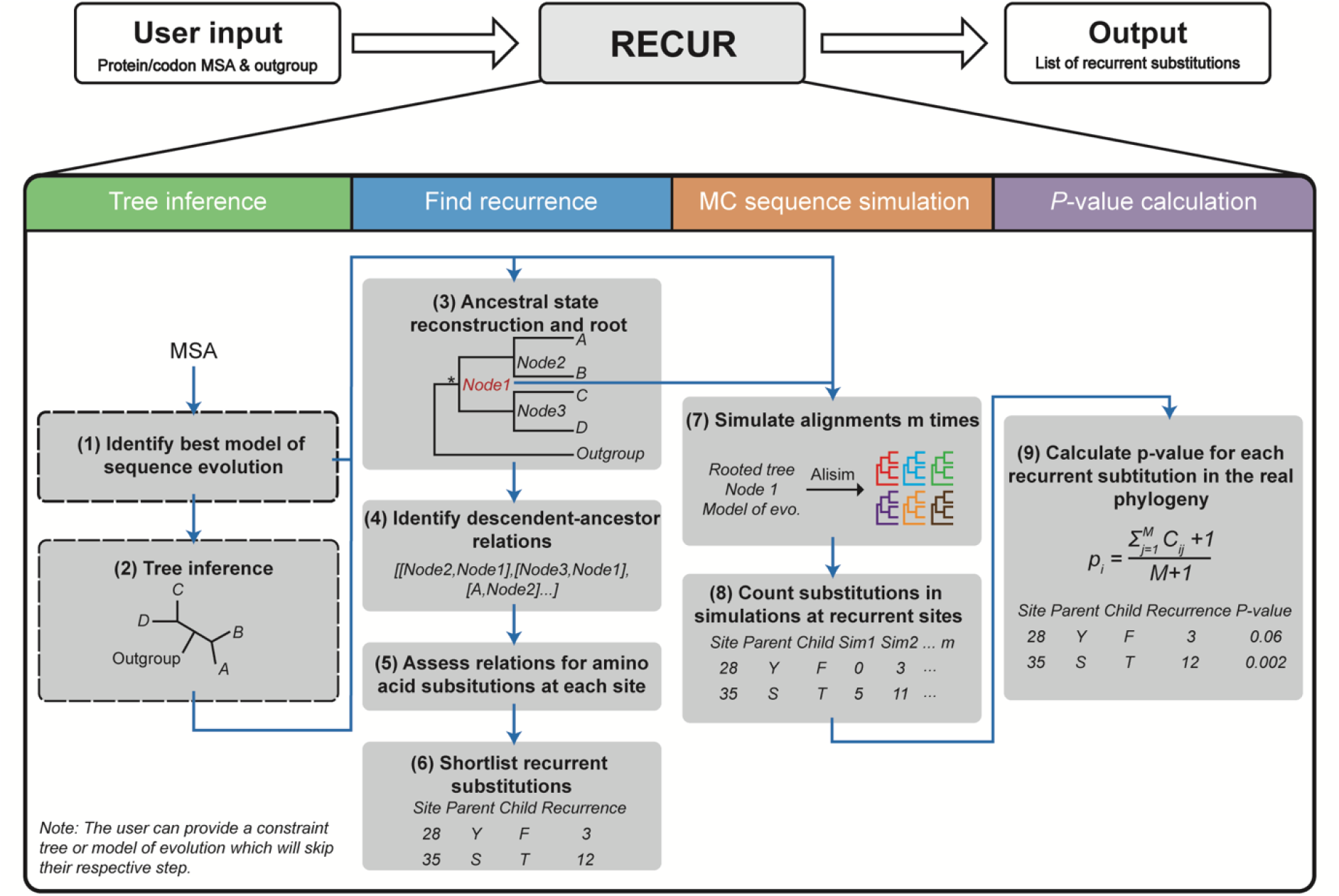
Overview of RECUR. The input (codon or protein) multiple sequence alignment is used to identify the best model of evolution (1), build a maximum likelihood phylogeny (2) and construct ancestral sequences (3). A user-defined outgroup is then used to root the tree (3) and assess each branch in the phylogeny for amino acid substitutions at each site in the protein sequence (4 and 5), from which recurrent substitutions are extracted (6). Sequence evolution is then simulated *m* times (default 1000) using the topology of the inferred phylogeny, best model of evolution and root sequence of the subtree of interest (*) which excludes the outgroup sequences (7). The recurrence of substitutions in the list output in (6) is then assessed in each of the simulated alignments (8) and a *p*-value is calculated from the number of times the recurrence in simulations equals or exceeds that observed in the real alignment (9).

### RECUR is fast and readily scalable to thousands of sequences

To demonstrate the runtime characteristics of the method we evaluated RECUR’s runtime and memory usage across multiple sequence alignments ranging from 8 to 1024 sequences. Each alignment comprised a randomly selected set of non-redundant SARS-CoV-2 surface glycoprotein (S protein) sequences (see Methods). Furthermore, to demonstrate the runtime characteristics of different steps of the method each alignment was subject to analysis with RECUR using eight threads with multiple different input options: 1) with no additional information, 2) with a pre-computed constraint tree, 3) with a pre-computed best-fitting model of sequence evolution, and 4) with both a pre-computed constraint tree and a pre-computed best-fitting model of sequence evolution.

The runtime is almost entirely consumed by tree inference (Figure 3A), and thus RECUR was fastest when a constraint tree was provided, as this enables RECUR to skip this step. In contrast, the memory usage (assessed as proportional set size, PSS) of the internal RECUR algorithm, which excludes tree inference, increased linearly with the number of sequences analysed (Figure 3B). Furthermore, only when >250 sequences were analysed did tree inference significantly increase memory consumption.

**Figure 3.**
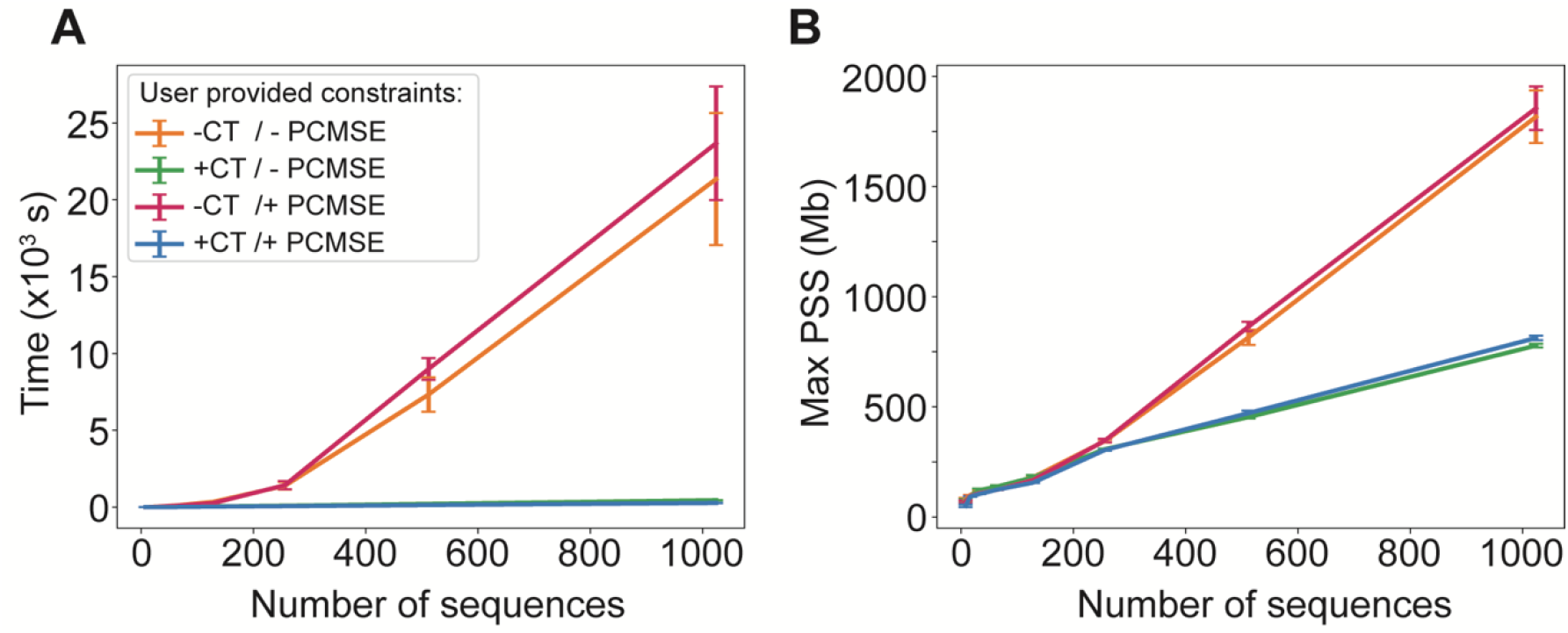
Analysing runtime and memory usage characteristics of RECUR. **A**) Line graph showing the relationships between runtime and number of sequences in the protein multiple sequence alignment. The different lines indicate different input variations; orange, no pre-computed constraint tree and no pre-defined model of sequence evolution (-CT/-PCMSE); pink, a pre-defined model of sequence evolution but no pre-computed constraint tree (-CT/+PCMSE); green, a pre-computed constraint tree but no pre-defined model of sequence evolution (+CT/-PCMSE) and blue, a pre-computed constraint tree and a pre-defined model of sequence evolution (+CT/+PCMSE). **B**) Line graph showing the relationship between maximum proportional set size (PSS) with the number of sequences. Lines coloured as in **A**.

### RECUR identifies widespread molecular recurrence during the evolution of the surface glycoprotein in SARS-CoV-2

To demonstrate the utility of RECUR, we applied the method to the complete set of non-redundant S protein sequences present in NCBI (n = 121,126, see Methods). The S protein, which forms a homotrimeric complex on the viral surface, is essential for host cell entry – mediating receptor recognition (predominantly the human angiotensin-converting enzyme 2, hACE2) via the exposed S1 subunit and facilitating membrane fusion through the more buried S2 subunit (Jackson, et al. 2022). This sequence was chosen as an illustrative example due to: (1) the clinical interest in the evolution of this protein sequence as it is the target of many vaccines; (2) the surface location and biological role of the protein means that the sequence has been subject to selection for promoting transmissibility and immune system evasion (Carabelli, et al. 2023) and thus is likely to have experienced substantial recurrent evolution (Korber, et al. 2020; Wilkinson, et al. 2022); and (3) the abundance of sequence data readily available in NCBI which can be subject to analysis. Application of RECUR to this dataset of S protein sequences identified 118,073 amino acid substitutions distributed across 98% (1,233/1,258) of aligned sites, with only 25 sites being invariant across all sequences in the alignment (Figure 4A and Supplementary File S1). A heatmap showing the frequency of the different types of amino acid substitutions inferred by RECUR is shown in Figure 4B. Of the 118,073 substitutions, 97% (114,890/118,073) were recurrent and parallel and involved 5,878 distinct substitution patterns (Supplementary File S2). The locations of these recurrent parallel substitutions were mapped onto the domains of the S protein (Figure 5A and B) revealing a landscape of widespread recurrent evolution along the length of the protein.

**Figure 4.**
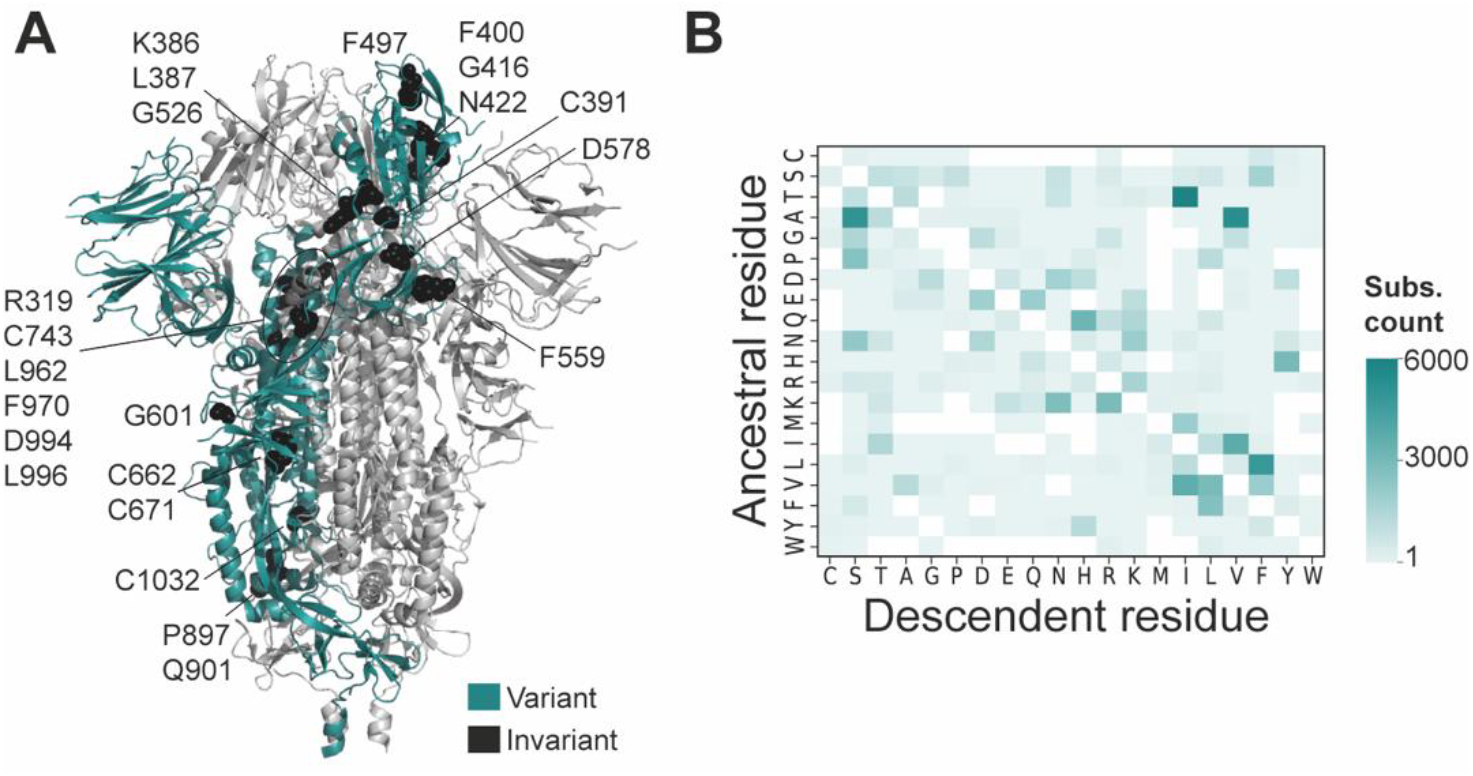
Position and type of amino acid substitutions inferred to have occurred by RECUR during the evolution of the S protein. **A**) Structure of the closed S protein trimer (PDB: 6VXX) with variable residues (where amino acid substitutions have been inferred) coloured cyan and invariant residues (no inferred amino acid substitutions) shown as black spheres on a single chain. Labels and approximate positions for the invariant residues are provided. Two sites (633 and 974) with no inferred amino acid substitutions but had site deletions (not reported by RECUR), are omitted. The remaining two protomers of the trimer are coloured in light grey. **B**) Heatmap of all amino acid substitutions identified by RECUR to have occurred in the S protein alignment across all sites. Rows signify the identity of the ancestral residue and columns signify the identity of the descendent residue.

**Figure 5.**
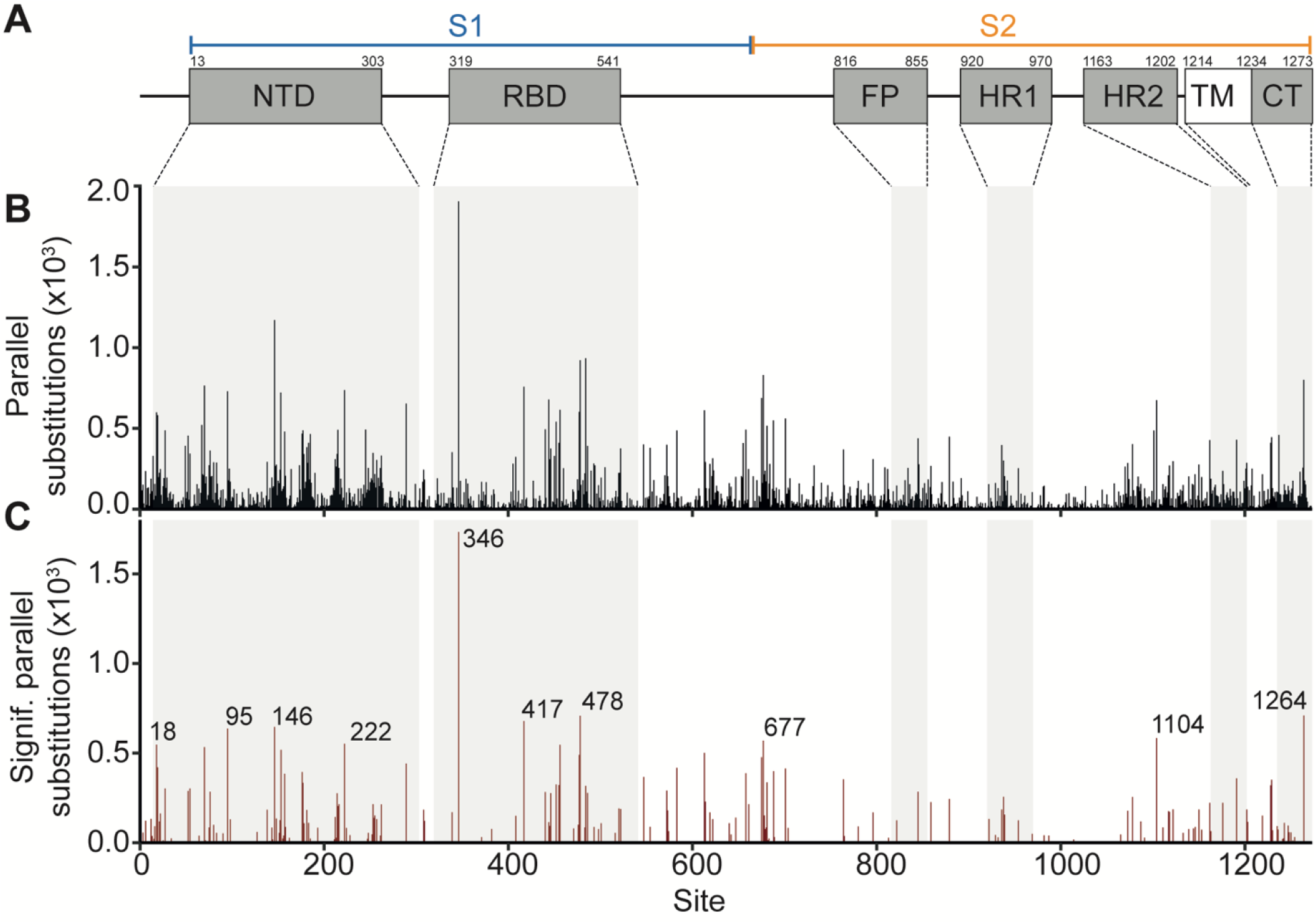
Recurrent evolution in the SARS-CoV-2 surface glycoprotein identified by RECUR. **A**) Map showing positions of the S protein domains (UniProt accession: P0DTC2). Domains include N-terminal domain (NTD), receptor binding domain (RBD), fusion peptide (FP), heptapeptide repeat sequence 1 (HR1), heptapeptide repeat sequence 2 (HR2), transmembrane domain (TM) and cytoplasmic tail (CT). The position of the larger S1 (residues 14 - 685) and S2 (residues 686 - 1273) subunits are indicated. **B** and **C**) Bar charts showing the number of parallel substitutions and significant parallel substitutions (those not likely the result of stochastic mutation) at each site in S protein, respectively. Protein domains are highlighted. The top ten sites with the most significant recurrent substitutions are labelled in **C**.

To determine whether these recurrent substitutions have occurred more frequently than expected given the number of sequences, the model of evolution, and the underlying phylogenetic relationship between the sequences, a Monte Carlo simulation of sequence evolution was performed. This revealed that 29% (33,596/114,890) of the recurrent substitutions identified above, which occurred at 206 different sites and included 356 distinct patterns of substitution, occurred significantly more frequently than expected (Figure 5C and Supplementary File S3). Among these sites, the top 10 with the most recurrent significant substitutions are highlighted in Figure 5C and Table 1. Eight of these sites are located in the exposed S1 region of the S protein, and include L18, T95, H146 and A222 in the N-terminal domain and R346, K417 and T478 in the receptor binding domain. All of these sites have all been associated with substitutions altering viral infectivity or immune escape (Table 1). For example, the L18F substitution (which occurred 453 times) is known to reduce neutralization for some antibodies (Harvey, et al. 2021; McCallum, De Marco, et al. 2021). Interestingly, eight of these sites experienced significant reversion substitutions (Table 1), indicating sequence space of the S1 protein is being resampled – a phenomenon associated with cycles of selection pressures influencing viral protein evolution (Focosi, et al. 2024). This pattern of reversion suggests that the S1 protein may be evolving within a constrained solution space, in which only a limited set of amino acid configurations are tolerated or advantageous under changing selective regimes. A wider examination of all significant recurrent substitutions revealed that 24% of significant recurrent substitutions (43 substitution pairs totalling 86 distinct substitution patterns) had a corresponding significant reversion substitution (Supplementary File S3). In summary, RECUR detected widespread recurrent molecular evolution during the evolution of the S protein including an abundance of reversion substitutions.

**Table 1.** The top 10 sites in the S protein with the most significantly recurrent parallel amino acid substitutions.

| Site | Region | Protein domain | Recurrent substitutions | Site annotation | Refs. |
| --- | --- | --- | --- | --- | --- |
| L18 | S1 | NTD | L→F<br>F→L | • In the NTD supersite for antibody binding.<br>• Mutations associated with immune escape. | (Grabowski, et al. 2021; McCallum, De Marco, et al. 2021) |
| T95 | S1 | NTD | T→I<br>I→T | • Mutations associated with immune escape and increased viral load. | (Shen, et al. 2021) |
| H146 | S1 | NTD | H→K<br>Q→K<br>K→Q | • In the NTD supersite for antibody binding.<br>• Mutations associated with immune escape. | (McCallum, De Marco, et al. 2021; Wang, et al. 2023) |
| A222 | S1 | NTD | A→V | • Mutations can alter ACE2 receptor binding (allosterically). | (Ginex, et al. 2022) |
| R346 | S1 | RBD | K→R<br>R→K<br>R→T<br>T→S<br>T→I<br>T→R | • Mutations associated with immune escape.<br>• Mutations can alter ACE2 binding affinity. | (Li, et al. 2024; Wang, et al. 2024) |
| K417 | S1 | RBD | N→K<br>K→N<br>N→T<br>T→K | • ACE2 receptor binding site.<br>• Mutations associated with immune escape. | (Barton, et al. 2021; Gupta, et al. 2024) |
| T478 | S1 | RBD | K→R<br>R→K<br>K→T<br>K→I<br>N→I | • Mutations associated with immune escape. | (Faraone, et al. 2023) |
| Q677 | S1 | - | Q→H | • Mutations can alter membrane fusion capability.<br>• Mutations can alter furin protease activity given proximity to cleavage site 682-RRAR-685. | (Zeng, et al. 2021) |
| V1104 | S2 | - | V→L<br>L→V | • Mutations may affect protein stability.<br>• Present in a T-cell epitope. | (Li, et al. 2024) |
| V1264 | S2 | CT | V→L<br>L→V | • In the 4.1, Ezrin, Radixin, Moesin binding domain. Thus, may affect subcellular trafficking of S protein. | (Hu, et al. 2023) |

### There is a significant overlap between significantly recurrent sites and those evolving under positive selection

Positive selection analyses have been extensively used to detect signatures of adaptive evolution by comparing synonymous and non-synonymous substitutions rates at individual sites. While this approach captures a distinct adaptive signature compared to that identified by RECUR, we sought to assess the degree of overlap between these two approaches. To do so, we compared sites identified as significantly recurrent by RECUR with those identified as evolving under positive selection based on multiple lines of evidence (Ferreira, et al. 2024). We found that 71% of sites identified as evolving under positive selection were also identified by RECUR as exhibiting significant recurrent substitution, demonstrating a significant overlap between the two distinct analyses (hypergeometric test, *p* < 0.001; Figure 6A and Supplementary File S3). This comparison suggests that while the two approaches identify distinct signals of adaptive evolution, an enriched subset of sites exhibit both significant recurrence and elevated rates of non-synonymous substitution.

**Figure 6.**
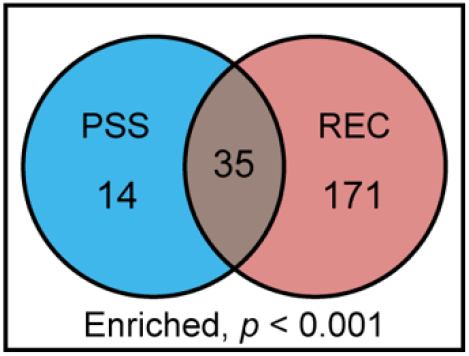
A Venn diagram showing the results of a hypergeometric test between sites under positive selection (PSS) (Ferreira, et al. 2024) and those identified by RECUR as having significant recurrent evolution (REC).

### Recurrent molecular evolution is enriched in the immune-exposed S1 subunit of the S protein

We next sought to investigate the position within the S protein of the significant recurrent substitutions identified by RECUR. This revealed a significant enrichment in the number of sites that have undergone recurrent evolution within the S1 subunit, the region which mediates target recognition and receptor binding and is also the primary target of the immune system (hypergeometric test, *p* < 0.01; Figure 7A). In contrast, there was a significant depletion in the number of recurrent sites located in the S2 subunit (hypergeometric test, *p* < 0.01; Figure 7B), the region that facilitates membrane fusion and endosomal trafficking. In agreement with these results and the S1 subunits increased exposure to the environment, sites that had undergone recurrent evolution had a significantly higher solvent accessibility compared to non-recurrently evolving sites when mapped onto the closed structure of the trimeric S protein (Wilcoxon rank sum test, *p* < 0.001; Figure 7C). These findings support that the S1 subunit, being more exposed on the viral surface making it more accessible to the host immune system, has experienced stronger selective pressures compared to the S2 subunit.

**Figure 7.**
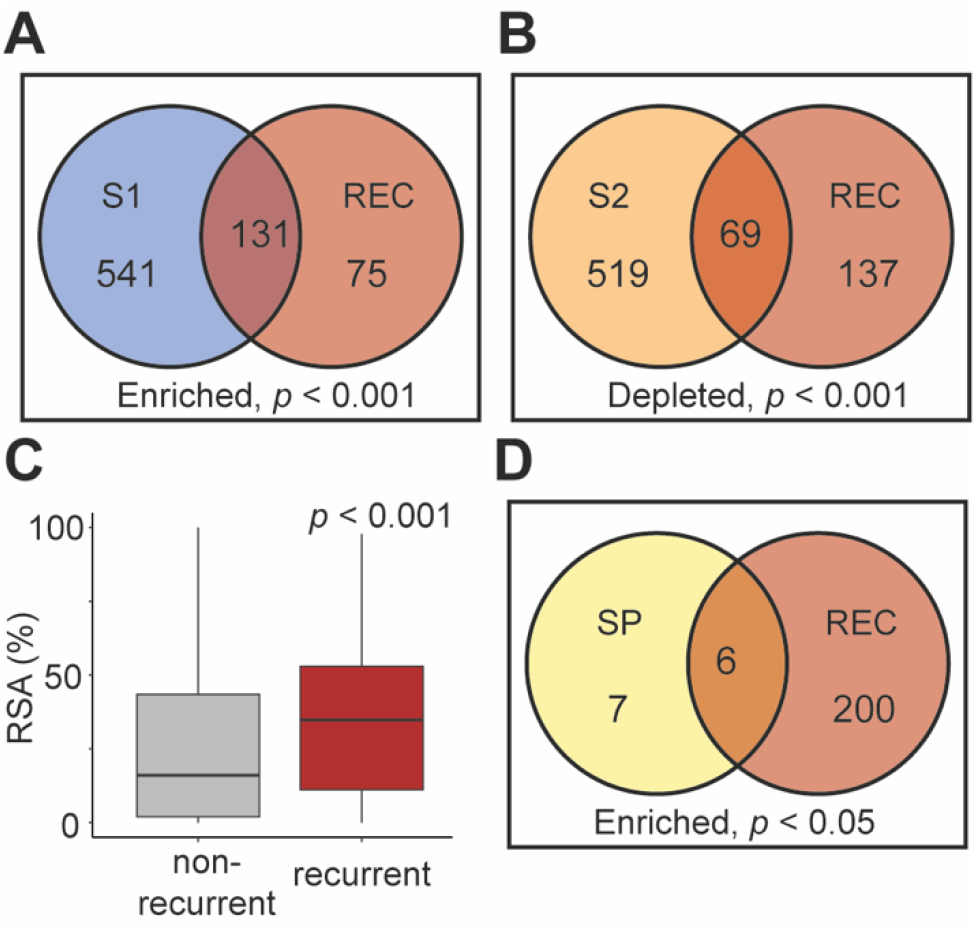
Assessing enrichment of recurrent evolution across regions of the S protein. **A and B**) Venn diagrams showing the results of hypergeometric tests between the S1 and S2 subunits with significant recurrence sites (REC), respectively. **C**) Boxplot showing the difference in the relative solvent accessibility (RSA) for sites identified as under significant recurrent evolution versus those that are not. The *p*-value indicates the significance of a Wilcoxon test. **D**) A Venn diagram showing the results of hypergeometric test between the signal peptide (SP) and significant recurrence sites.

For completeness we also investigated the extent of recurrent evolution in the signal peptide, the N-terminal 13 amino acids upstream of the S1 region responsible for entry into the host’s endocytic system. This revealed an enrichment of significant recurrent evolution within the signal peptide (hypergeometric test, *p* < 0.05; Figure 7D). One of the recurrent substitutions identified in the signal peptide was S13I which was inferred to have 37 occurrences across the phylogeny (Supplementary File S3). The S13I substitution has been shown to shift the signal peptide cleavage site leading to structural alterations to the N-terminal domain interfering with the binding of some antibodies (McCallum, Bassi, et al. 2021). Furthermore, S13I may increase the efficiency of protein secretion (Zhang, et al. 2022). Thus, although often overlooked, the signal peptide of the S protein has hallmarks of having undergone adaptive evolution, with some substitutions already shown to influence viral replication and immune evasion.

### Recurrent molecular evolution has been restricted at the trimeric interface but enhanced at the hACE2 interface of the S protein

To further investigate the functional implications of recurrent substitutions identified by RECUR, we next examined the protein-protein interfaces of the S protein, focusing on the pre-fusion trimeric interface and the human angiotensin-converting enzyme 2 (hACE2) receptor binding interface. To do this, residues at the trimeric and hACE2 interfaces were first identified using a distance threshold of less than 4 Å to an atom of the respective neighbouring protein (Figure 8A and B, Supplementary Files S4). An analysis of these sites revealed that there was a depletion of sites that have undergone significant recurrent evolution at the pre-fusion trimeric interface of the S protein complex (hypergeometric test, *p* < 0.01; Figure 8C). In contrast, recurrently evolving sites were enriched at the interface with the hACE2 receptor (hypergeometric test, *p* < 0.01; Figure 8D). Thus, recurrent evolution has been constrained at the trimeric interface, likely reflecting functional or structural limitations, whereas recurrent evolution is enriched at the immune-accessible hACE2 binding site of the S protein, where adaptive pressures such as host immune evasion drive repeated amino acid changes.

**Figure 8.**
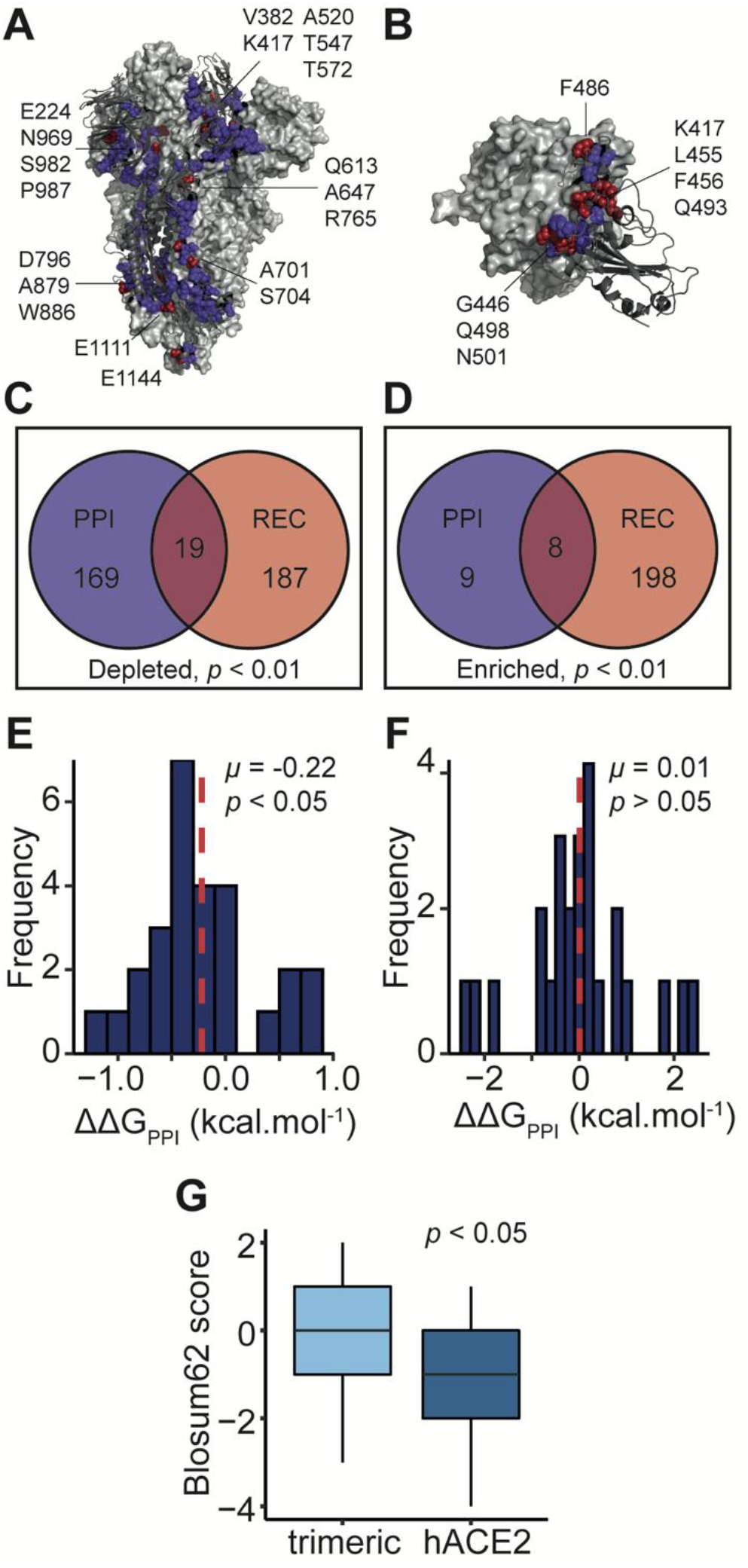
The effect of recurrent substitutions at protein-protein interfaces formed by the S protein. **A**) The trimeric interface of the S protein complex (PDB: 6VXX). A single S protomer is shown as a dark grey cartoon. The residues interacting with the remaining two protomers (the light grey surface) are shown as spheres. Interacting residues that have undergone significant recurrent evolution are shown in red and the residue labelled. **B**) Interface between the S protein RBD (dark grey cartoon) and the human angiotensin converting-enzyme 2 (hACE2; light grey surface) (PDB: 6M0J). Interface residues are shown as in **A. C** and **D**) Venn diagrams showing the results of hypergeometric tests between the protein-protein interface (PPI) residues with significant recurrence sites for the trimeric and hACE2 interfaces, respectively. **E** and **F**) Histograms showing the effect of recurrent substitutions on the interaction energy of their respective interface (ΔΔG_PPI_). The mean value (*μ*) is indicated with a dashed red line and *p*-values are the results of one sample t-tests. Bin width is 0.2 kcal.mol^-1^. **G**) Boxplot showing the difference in Blosum62 score between recurrent substitutions occurring at the trimeric interface versus the hACE2 interface. The *p*-value of a two-sample t-test is shown.

### Recurrent molecular evolution has stabilised the trimeric S-protein interface and modulated receptor binding and immune evasion at the hACE2 interface

Finally, we assessed the impact of the significant recurrent substitutions on both the trimeric pre-fusion and hACE2 interfaces. To do this, we calculated the change in the interaction energy (ΔΔG_PPI_) upon *in silico* mutation (see Methods), where a negative ΔΔG_PPI_ indicates interface stabilization and *vice versa*. Our analysis revealed that recurrent substitutions at the trimeric interface predominantly act to stabilize the interaction (mean = -0.22 kcal.mol^-1^, one sample t-test *p* < 0.05; Figure 8E). Furthermore, two of these substitutions (K969N and V987F) had a |ΔΔG_PPI_| > 1 kcal.mol^-1^ and were both predicted to stabilize the pre-fusion trimeric interface. Meanwhile, the recurrent substitutions at the hACE2 interface had no overall effect on stability (mean = 0.01 kcal.mol^-1^, one-sample t-test *p* > 0.05; Figure 8F). Of these substitutions, seven had a |ΔΔG_PPI_| > 1 kcal.mol^-1^, with substitutions away from the reference sequence (F486P, F486V and K417N) destabilising the hACE2 interface and reversions back to the reference sequence (P486F, T417K, V486F and N417K) stabilising the hACE2 interface. Notably, all three hACE2 destabilising substitutions have been associated with immune escape (Cao, et al. 2022; McCallum, et al. 2022; Qu, et al. 2024). In line with the above results, the variance in ΔΔG_PPI_ was significantly greater at the hACE2 interface in comparison to the trimeric pre-fusion interface of the S protein (Levene’s test, *p* < 0.01). Lastly, the substitutions occurring at the hACE2 receptor interface were significantly less conservative than those occurring at the pre-fusion trimeric interface (two-sample t-test, *p* < 0.05; Figure 8G). Together, these results show that recurrent substitutions have primarily acted to stabilize the S protein’s trimeric interface, whereas their effects at the hACE2 binding interface are more variable. The latter supports the notion that adaptive evolution at the hACE2 interface is balancing immune system evasion with maintaining hACE2 binding capabilities.

## Discussion

Recurrent evolution is a pervasive feature across the tree of life and a hallmark of adaptive evolution. At the molecular level, recurrent sequence changes provide powerful insights into the selective pressures shaping protein evolution. However, detecting such patterns at scale remains challenging. Here, we present RECUR, a scalable tool that automatically detects recurrent amino acid substitutions from a protein (or an equivalent codon) multiple sequence alignment. RECUR enables the quantification of recurrent evolution across large datasets, revealing which amino acid states are repeatedly favoured by natural selection. We applied RECUR to 123,126 surface glycoprotein sequences from SARS-CoV-2, identifying key sites that have undergone recurrent evolution during viral adaptation. This tool opens up new avenues for investigating the molecular basis of adaptation and constraint across diverse biological systems.

To demonstrate RECUR’s capabilities, we applied it to the complete set of non-redundant SARS-CoV-2 surface glycoprotein sequences derived from human hosts, revealing widespread recurrent evolution across the length of the protein. Furthermore, we uncovered a significant enrichment of sites that have undergone recurrent evolution in the S1 subunit compared to a significant depletion in the S2 subunit. This aligns with the functional and structural differences between these two regions of the S protein. The S1 subunit, contains the receptor binding domain and is essential for viral attachment to the hACE2 receptor, making it a hotspot for adaptive evolution as the virus evolves to concurrently optimise binding affinity and evade the immune system (Tian, et al. 2021). Given the trade-offs between these factors there are high levels of recurrent reversion substitutions in this region of the S protein as the virus undergoes adaption during cycles of selection pressure. Additionally, the S1 subunit contains the N-terminal domain, which contributes to antigenicity and hence adaptative evolution has repeatedly evolved mechanisms for immune escape in this region (McCallum, De Marco, et al. 2021). Finally, although depleted, sites under recurrent evolution were also identified in the S2 subunit. These substitutions may benefit the virus through aiding immune escape through allosteric effects on S1 epitopes (Kumar, et al. 2023). Importantly, this analysis has identified many sequence changes for which we could find no existing experimental interrogation, but which have repeatedly evolved during the radiation of SARS-CoV-2. Future functional interrogation of these sites may reveal novel mechanisms of viral adaptation and uncover previously unappreciated determinants of viral fitness. While our analysis is not intended to be exhaustive, it highlights RECUR’s utility as a tool for the identification of key residues under recurrent evolution, providing a framework for prioritising specific residues and substitutions for downstream evolutionary and functional studies.

To support these applications, we evaluated RECUR’s runtime characteristics. The runtime of RECUR is influenced by the number of sequences, the sequence length, and the extent of recurrence present in the sequences under consideration. Larger numbers of sequences, longer alignments, and higher rates of recurrence each result in increased computational demand. The SARS-CoV-2 surface glycoprotein chosen to exemplify the method has a high mutation rate, is substantially longer than the average eukaryotic or bacterial protein (S protein length = ∼1300, average prokaryotic protein length = ∼300, average eukaryotic protein length = ∼500 (Tiessen, et al. 2012)), thus runtimes presented in this study represent a conservative estimate of what can be expected by a user when running RECUR. Furthermore, our benchmarking revealed that most of RECUR’s runtime is consumed by the phylogenetic tree inference step, not by the recurrence detection algorithm itself. To support more flexible analyses, RECUR allows users to input a pre-computed constraint tree forcing RECUR to use a fixed topology. This provides the benefit of leveraging more information to infer the constraint tree than may be available in the alignment under consideration. For example, the user may wish to constrain the topology of the tree to match a known species tree when analysing orthologous sequences derived from those species.

The RECUR method makes extensive use of the IQ-TREE 2 software package (Minh, et al. 2020). IQ-TREE 2 was chosen over other phylogenetic inference software for several reasons. First, IQ-TREE 2 is a mature, well-developed and thoroughly benchmarked software package that has the capability to carry out many steps required in the RECUR algorithms. These steps include model selection, tree inference, ancestral state reconstruction, and alignment simulation. It is not possible to achieve all of these steps using comparable software such as RAxML (Stamatakis 2014) or PhyML (Guindon and Gascuel 2003). Thus, use of IQ-TREE 2 reduces the number of dependencies required by RECUR. Secondly, IQ-TREE performs very well against all competitor methods in terms of both speed and accuracy (Zhou, et al. 2017). Thirdly, IQ-TREE 2 can analyse alignments with thousands of sequences allowing RECUR to also scale to this number of sequences. Together, these features ensure RECUR is both robust and scalable.

It is important to note that RECUR is not intended to identify positive selection. Traditional positive selection approaches detect adaptation by comparing rates of synonymous and non-synonymous substitutions, identifying sites where the rate of non-synonymous substitution is higher than expected. In contrast, RECUR focuses on the recurrence of substitutions, identifying specific substitutions that have higher levels of recurrence than expected given the number of sequences sampled, the phylogenetic relationship of those sequences, and the best fitting model of sequence evolution identified from the input alignment. That being said, there was significant overlap in the sites detected by both approaches, suggesting that many adaptively evolving sites exhibit multiple signals of adaptive evolution. Meanwhile, the differences of sites between methods reflect their unique detection criteria. Given these findings, RECUR can serve as a valuable tool for studying adaptive evolution, either as a standalone approach or in conjunction with positive selection analyses to provide additional insight into the evolutionary dynamics of protein sequences. Furthermore, while RECUR uses a single transition matrix for all sites in the protein sequence, it has been previously shown that sites evolving under strong purifying selection are typically depleted for recurrent substitutions (Robbins and Kelly 2024). This suggests that the recurrence signal is unlikely to arise solely from limited amino acid tolerability at a site, but rather repeated adaptive change.

In conclusion, RECUR is a comprehensive tool for identifying amino acid substitutions from multiple sequence alignments. We anticipate that RECUR will serve as a valuable resource for understanding the molecular basis of convergent phenotypes as well as generating hypotheses and guiding experimental testing of protein function.

## Materials and methods

### Implementation

RECUR is a Python application that leverages IQ-TREE 2 (Minh, et al. 2020) as an external dependency and requires Python 3.9 or higher. The application utilizes DendroPy (Sukumaran and Holder 2010) to extract ancestor-descendent sequence pairs from unrooted trees in Newick format. RECUR achieves high performance through the use of NumPy (Harris, et al. 2020) and multiprocessing, particularly in key steps such as the substitution analysis of Monte Carlo simulated protein sequences. In this process, the ancestor-descendent sequence pairs are first converted into numerical representations and then into NumPy arrays for element-wise comparison at each residue site. Multiprocessing is employed to further speed up these operations. RECUR includes a Linux version of the IQ-TREE 2 binary, enabling it to run as a standalone application on Linux systems. For Windows and macOS users, IQ-TREE 2 must be preinstalled. Regardless of the operating system, it is recommended to run RECUR within a conda environment or a Docker container for compatibility. Full instructions on the installation and implementation of RECUR can be found at https://github.com/OrthoFinder/RECUR.

### Tree inference and model selection

RECUR takes a codon or protein multiple sequence alignment as input. The alignment is evaluated to identify the best-fitting model of sequence evolution using ModelFinder implemented in IQ-TREE 2 (Kalyaanamoorthy, et al. 2017). Using this model, a maximum likelihood phylogenetic tree is built using IQ-TREE 2’s ultrafast bootstrapping method with 1000 replicates (Hoang, et al. 2017). Alternatively, if the user has already pre-computed a phylogenetic tree or pre-selected a model of sequence evolution, then these can be provided as inputs to RECUR and be used in all analysis steps. The tree is then rooted on a user-defined outgroup which can comprise a single sequence or a list of outgroup sequences. In the case where the supplied outgroup sequences are not monophyletic in the tree, the smallest possible clade comprising the full set of outgroup sequences will be selected as the outgroup.

### Identifying recurrent amino acid substitutions

Ancestral sequence reconstructions are inferred for every node in the phylogeny using the best-fitting model of sequence evolution and the inferred maximum likelihood tree using the ancestral state reconstruction method implemented in IQ-TREE 2 (Minh, et al. 2020). To correctly infer gaps in ancestral sequences (which IQ-TREE 2 currently cannot implement) the multiple sequence alignment is converted into binary sequences, with 0 and 1 representing gapped and non-gapped sites, respectively. As with the biological sequences, the ancestral reconstructions of the binary sequences are inferred for every node in the phylogeny using the inferred maximum likelihood tree and the general time reversible model for binary data model (GTR2) which allows for unequal state frequencies. The positions of the inferred 0 values, i.e., gaps, in the ancestral sequences are then mapped onto the biological sequences to complete ancestral sequence reconstruction with indel estimation.

If a codon alignment was provided, both the inferred and extant sequences are then translated into protein sequences using the standard genetic code as default. The user can also specify alternative NCBI genetic codes to translate DNA sequences if required. For each residue in the protein sequence, amino acid substitutions are tabulated by analysing the residue identity for all ancestor-descendent sequences at every branch in the subtree that excludes outgroup sequences. A list of recurrent substitutions, i.e., substitutions that occurred more than once at a given site during the evolution of the sequence set, is then compiled.

### Identification of significant recurrent sites

The phylogeny, model of evolution, and root sequence of the subtree of interest are then used to simulate codon/protein sequence evolution *M* times (default 1000) using the alisim function in IQ-TREE 2 (Ly-Trong, et al. 2023). As explained above, the gaps are mapped onto each simulated alignment. This prevents the overestimation of substitution recurrence. The amino acid substitutions that have occurred in each of the simulated sequence alignments are then identified using the method outlined above. For each of the recurrent substitutions identified for the real alignment, RECUR assesses the number of times that site specific substitution has occurred at the same or greater frequency in the simulated alignments. A *p*-value describing the probability that the recurrence of that site specific amino acid substitution having occurred by chance (*p*_i_) is then assigned using the following equation:

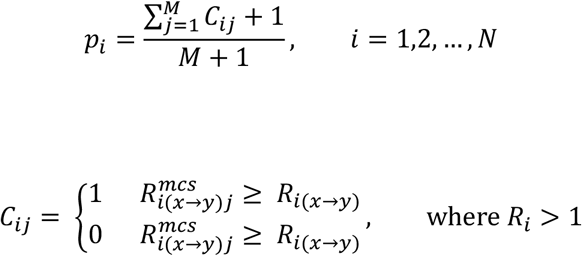

where *i* is the *i*th residue, *N* is the number of residues in the protein alignment, *j* is *j*th sequence simulation, *M* is the number of sequence simulations run, *C*_*ij*_ is the piecewise function, *R*_*i(x→y)*_ is the recurrence of amino acid substitution *x → y* at the *i*th site in the real phylogeny, and *R*^*mcs*^_*i(x→y)j*_ is the recurrence of amino acid substitution *x → y* at the *i*th site in the *j*th simulation.

### Data set retrieval and curation

A dataset comprising all available SARS-CoV-2 surface glycoprotein coding sequences derived from human hosts were downloaded from National Center for Biotechnology Information on 29^th^ July 2024 (n = 3,934,602). The dataset was filtered to remove sequences shorter than 3,800 bp, sequences with ambiguous characters and sequences without start or stop codons. The coding sequences were then translated into protein using the standard genetic code and aligned using FAMSA (Deorowicz, et al. 2016). The alignment was trimmed to remove columns with >50 % gaps and non-redundant sequences were removed resulting in an alignment containing 123,126 SARS-CoV-2 surface glycoprotein sequences with 1273 columns. Finally, 15 sites known to cause problems with phylogenetic analyses were obtained from https://genome.ucsc.edu/cgi-bin/hgTables and masked in the alignment. The final alignment is provided in Supplementary File S5. The sequence identical to the Wuhan-1 SARS-CoV-2 surface glycoprotein (accession: MT582461.1) was used as the outgroup.

### Computing runtime and memory usage

To demonstrate the runtime characteristics, the 123,126 sequence multiple sequence alignment was randomly sampled, while preserving the outgroup species, to create 10 alignments with 8, 16, 32, 64, 128, 256, 512 and 1024 sequences. RECUR was run on each alignment with four input option combinations: no constraint tree and no evolutionary model, a constraint tree but no evolutionary model, an evolutionary model but no constraint tree, and both a constraint tree and evolutionary model. The evolutionary model and constraint trees used were those obtained from the RECUR output under the first input scenario. Each run was timed, and maximum proportional set size (PSS) was recorded. These tests were conducted on an AMD EPYC 7742 64-Core Processor (2.25 GHz, x86_64 architecture) and given eight threads.

### Analysing the complete 123,126 surface glycoprotein sequence dataset

IQ-TREE 2 is unable to infer phylogenetic tree from an alignment comprising 123,126 sequences. Thus, a maximum likelihood constraint tree was inferred using Very Fast Tree (Piñeiro, et al. 2020), which is specifically designed for large datasets. In addition, the best fitting model of sequence evolution and the averaged model parameters, HIVw+F+I[0.24]+G4[0.59], was taken from the runtime and memory usage runs for the 1024 sequence alignment runs above. By constraining the tree and the model of sequence evolution RECUR is thus able to analyse the complete dataset.

### Residue solvent accessibility analysis

The relative solvent accessibility (RSA%) was determined for each residue in the closed S protein trimeric structure (PDB: 6VXX) using the GetArea web server (https://curie.utmb.edu/getarea.html) (Supplementary File S6).

### Protein-protein interface analysis

Protein-protein interface residues were identified using the EMPL-EBI PDBsum database. To analyse the trimeric S protein interface, the closed (PDB: 6VXX) and open (PDB: 7W92) structures were utilized, while the hACE2 interface was examined using the 6M0J structure (Supplementary File S4). The effects of recurrent substitutions were assessed by introducing single amino acid substitutions using PyMol. Then, the change in Gibb’s free energy of interaction (ΔG_PPI_) was calculated using BAlaS at the respective interface for both the wild-type and mutated structure (Wood, et al. 2020). From these, the change in ΔG_PPI_ can be calculated, where a negative value indicates a strengthening of the interface, and a positive value indicates a weakening of the interface (Supplementary File S3).

## Acknowledgements

This work was funded by the Wellcome Trust under grant agreement number 226598/Z/22/Z. ER is funded by the Wellcome Trust and the Biotechnology and Biological Sciences Research Council (BBSRC) [grant numbers BB/M011224/1 and BB/P003117/1]. This research was funded in whole, or in part, by the BBSRC. For the purpose of open access, the author has applied a CC BY public copyright license to any Author Accepted Manuscript version arising from this submission.

## Data Availability

The source code is available on the RECUR GitHub repository (https://github.com/OrthoFinder/RECUR). All data used in this study is provided in the supplementary material.

## Author Contributions

ER and SK conceived the study. ER and YL wrote the software. ER and YL carried out the analysis. ER and SK wrote the manuscript.

